# Systematic map of the most recent evidence (2010 to 2019) on ruminant production-limiting disease prevalence and associated mortality in Ethiopia

**DOI:** 10.1101/2022.07.15.500201

**Authors:** Theodora K. Tsouloufi, Isla S. MacVicar, Louise M. Donnison, Karen L. Smyth, Andrew R. Peters

## Abstract

Ethiopia’s livestock sector supports the livelihoods of millions of smallholder farmers. However, despite the improvements of recent years, livestock productivity remains low due to critical constraints, including infectious diseases. The aim of this study was to collate and synthesise the published evidence on ruminant disease frequency and disease-associated mortality in Ethiopia, by identifying knowledge gaps and clusters in the literature to provide the basis for a decision-making tool. Searches on both bibliographic and organisation databases were conducted in English and were restricted to the period 2010-2019. Search results were screened for relevance at title, abstract and full text level, which identified 716 articles relevant to the research question. The systematic map revealed an increased publication output from 2012-2017, compared to 2010-2011 and 2018-2019. Most studies were conducted in Oromia, Amhara and SNNPR. A substantial body of evidence was found for trypanosomosis, ectoparasite infestation, fasciolosis, nematodiasis, echinococcosis, brucellosis and bovine brucellosis. This study suggests that despite the high output of epidemiological publications, further understanding of a considerable number of diseases is required and where evidence is abundant, synthesis of information should be carried out in order to better inform decisions on disease control priorities in the livestock sector.

## Introduction

Livestock are catalytic in supporting the livelihoods of millions of poor smallholders in sub-Saharan Africa, in both rural and peri-urban settings (Randolph *et al*., 2007). Ethiopia has the largest livestock population in Africa and the ninth largest population in the world. Ruminants account for 80% of the national herd and contribute significantly to poverty alleviation, with small ruminants being key to the improvement of the income status of female smallholder farmers (Shapiro *et al*., 2017). Despite the constantly growing ruminant population and the dairy sector’s potential over the last decade, the livestock subsector in Ethiopia is facing serious constraints, such as reduced livestock productivity, which impair its growth (Minten *et al*., 2020; Leta and Mesele, 2014). Infectious diseases are a highly ranked cause of compromised livestock productivity in the country, with disease-associated losses amounting to hundreds of millions of dollars in specific years (Berhanu, 2002). The absence of a clear policy framework for specific aspects of livestock development, including disease control, has been considered an additional hurdle to the sector’s growth (Legese and Fadiga, 2014). On these grounds, disease-associated livestock morbidity and mortality have become a highly relevant topic for international policy and research in Ethiopia, and a considerable number of research initiatives have been generated over recent years, resulting in an increasing volume of evidence. Although, there have been some comprehensive efforts to collate evidence for specific ruminant diseases in the country, no systematic mapping of the whole body of evidence has so far been undertaken. This significant knowledge gap thus poses a considerable challenge for decision-makers who strive to define the problem and establish its magnitude.

Systematic evidence mapping is a methodology extensively used in the social sciences and the environmental sector to provide an overview of the evidence landscape in a research area by exposing knowledge gaps and clusters (Wolffe *et al*., 2019). Although systematic mapping reviews share the rigorous approach of a systematic review, they usually answer research questions with a wider scope and do not typically involve full data extraction, validity assessment or quantitative synthesis (i.e. meta-analysis) of the research results (Dicks *et al*., 2014). Systematic maps can additionally include bibliometric data, which can be used to assess journal quality, worldwide research trends and priorities, and the dynamics of interdisciplinary collaborations. Researchers and decision-makers can use systematic maps to gain information about gaps or under-reported topics in the current evidence base, as well as to benefit from highlighted knowledge clusters that allow for a full systematic review synthesis. Researchers can also benefit from the identification of deficiencies and best practices across the evidence base, which may be used to increase consistency across studies (Haddaway *et al*., 2016). Most importantly, funders can appreciate research areas that have previously received a considerable amount of attention and can therefore, identify priorities and allocate accordingly their funds to future research (Haddaway *et al*., 2016).

Data gaps impact investments in the livestock sector and thus curtail its potential for future growth, economic development, and poverty alleviation. An estimated 25-40% of published literature on specific livestock diseases in sub-Saharan Africa is of poor quality and difficult to summarize due to the diversity of studies (Alonso *et al*., 2016). Improvements in livestock health data accessibility and quality have therefore, been identified as key actions to increase livestock productivity in Low- and Middle-income countries (LMICs). The systematic map methodology could be a promising decision-making tool for LMICs, as disparate evidence hampers evidence-based decision-making.

Considering the above, the objective of this systematic mapping review was to: i) collate and synthesise the published evidence on ruminant disease frequency and disease-associated mortality in Ethiopia, ii) identify knowledge gaps and clusters in the literature in order to inform future research and iii) provide the basis for a decision-making tool in the livestock sector.

### Stakeholder engagement

The topic for this systematic map arose from a Bill and Melinda Gates Foundation funded project, Supporting Evidence Based Interventions-Livestock (SEBI-Livestock), established to collate better livestock data to help inform decision-making. The scope of the project was then refined through expert discussions, which helped to formulate the research question. A diverse group of stakeholders were involved in discussions of the scope, protocol review, search strategy and final presentation of the map, all of whom were members of the Livestock Data for Decisions Community of Practice (https://www.livestockdata.org).

### Objectives of the map

The main research question of this systematic mapping review was:

“*What is the most recent available evidence on ruminant infectious disease frequency and disease-associated mortality in Ethiopia?”*

The key objective, as requested by the stakeholders, was to catalogue the existing evidence across several production-limiting ruminant infectious diseases and of disease-associated mortality. Moreover, although systematic maps do not typically capture study details, we chose also to extract the prevalence values for the respective diseases, as this was identified as important information for our stakeholders. The envisaged output of this process was to establish a comprehensive interactive evidence map of ruminant disease and mortality research in Ethiopia, along with the respective reported prevalence.

The research question framed according to the Population and Outcome (PO) scheme was:

- **Population**: Ruminants (cattle, sheep, and goats) reared in Ethiopia
- **Outcome**: Distribution of evidence, disease incidence/prevalence, disease-associated mortality

## Methods

### Registration of the protocol

The present review was conducted in accordance with the protocol previously published (Tsouloufi *et al*., 2020), which conformed to the Reporting Standards for Systematic Evidence Syntheses (ROSES) and PRISMA-ScR standards (Haddaway *et al*., 2017). Reporting standards such as ROSES and PRISMA are widely accepted in the evidence synthesis community to support transparency and rigor of the research (Haddaway and Macura, 2018).

### Deviations from the protocol

A few modifications to the original protocol were made in order to facilitate the review process. Although we stated in our protocol that we would consider an evaluation of the methodological quality of the included papers, we have not included such an analysis in this review due to time and resource constraints. This will be discussed extensively in the context of a future publication and is not considered a critical omission for this analysis, as the quality of included studies is not typically appraised in the context of a systematic map, according to the established methodological guidance (James *et al*., 2016). Critical appraisal in systematic maps is optional since there is no synthesis of the results and it is difficult to assess external validity when the research question is not explicitly specified as it is with a systematic review. Furthermore, although this was not intended to be a bibliometric study, we added a quality metric of the published studies according to their journal ranking, as we felt that this would give a preliminary insight to the quality of research produced. For this purpose, we opted for Scimago Journal & Country Rank for the scientific journals (https://scimagojr.com), as this has previously been used in systematic maps (Alla and Nafil, 2019). Finally, we excluded the “case definition” category from the data extraction form, as almost none of the included studies succinctly defined the suspected/confirmed case for the respective diseases, which thus precluded any further evaluation.

### Searches

A comprehensive search of multiple information sources attempted to capture an un-biased sample of literature to encompass both published and grey literature. Initially all searches were conducted within a period of 3 months (January-March 2020). In addition to these searches, the bibliographic database searches were repeated within the time period of a month (November 2021).

### Search terms

Boolean search strings in the form of (Population) AND Ethiopia AND (Outcome) were developed for each of the studied conditions according to the suggestions made by specialist librarians within the University of Edinburgh. The respective search strings were tested in the first instance in the selected databases and were accordingly adapted in reviewers’ meetings until finalized. Specifically, a benchmark of 5 highly relevant articles (Supplementary File 1) were screened against scoping search results to examine whether searches could locate relevant evidence. If the database did not allow for a full search string, a shortened version was produced and adapted as necessary. Searches were restricted on publication dates from January 2010 to December 2019. Details of the terms used in each of the searches are provided in Supplementary File 1, as there were variations between search terms employed across the different databases. The number of articles retrieved by each search term were recorded in the Supplementary File 1. The initial searches were performed in 2020. The bibliographic databases and the Addis Ababa University repository searches were repeated in 2021. Despite the number of search results being recorded in 2020, the associated endnote files were not archived before screening. In 2021, the searches were repeated in order to ensure that there were copies of the Endnote files archived and the process could be repeated. These repeated searches were also performed to generate metrics on the relative importance of each database to the final search results. These results are informing the prioritisation of ongoing work automating the process using machine learning techniques (Goldfarb-Tarrant *et al*., 2020). Supplementary File 1 details the results of both the searches in 2020, as well as the searches in 2021.

### Bibliographic databases

The following online bibliographic databases were searched to identify relevant literature: Scopus, Web of Science (Core collection), MEDLINE^©^ via NCBI^©^ and CAB direct^©^ via CABI^©^.

### Search engines and specialist websites

Grey literature was captured by search engine searches using Google Scholar (500 first results sorted by relevance) and GrayLit network. Additionally, the databases Networked Digital Library of Theses and Dissertations and the website African Journals Online were searched.

### Organisational websites

The following specialist websites were also searched:

International Livestock Research Institute (https://www.ilri.org), the Consultative Group for International Agricultural Research (CGIAR) (https://www.cgiar.org/), the Food and Agriculture Organization of the United Nations (FAO, http://www.fao.org), the United States Agency for International Development (USAID, https://www.usaid.gov/), World Organization for Animal Health (OIE, http://www.oie.int/) and, the electronic repository of the Addis Ababa University (http://etd.aau.edu.et/)

### Endnote files and compilation of search results

Results of the searches in bibliographic databases were downloaded as reference files and assembled as a final library using the desktop version of Endnote® X9 (Clarivate Analytics). Duplicates were automatically removed after all endnote files were combined. Results from the other sources (search engines and websites) were saved as screen captures and were categorised into separate files. Duplicates of these searches were manually removed prior to screening.

Records from bibliographic databases were introduced into the screening workflow. Records from other sources were screened separately before they were combined with other records. Due to the project’s restricted timelines, snowballing (i.e. use of paper’s reference list to identify additional papers) and update searches were not performed.

### Study screening

All studies retrieved from searches were screened via a two-stage screening process (title-abstract and full-text level) following the CEE guidelines (CEE, 2018), and by applying the pre-defined eligibility criteria. In the first stage, all titles and abstracts were prospectively evaluated by two independent reviewers (TKT, ISMV) in terms of their relevance to the research questions. Studies where the title and abstracts meet the eligibility criteria were evaluated in their full length for inclusion. Any doubt over the presence of a relevant inclusion criterion or where abstracts were lacking, resulted in the articles being retained for assessment at a full-text level. At the beginning of each stage, the agreement between the two reviewers (LMD and ISMV) was evaluated in a random subset of papers (1000 articles screened at both the title and abstract stage, as well as the full-text stage) with the use of the Cohen’s Kappa statistic and in order to ensure consistency and common understanding of the eligibility criteria. A Kappa score of 0.69 indicated substantial agreement for inclusion between the reviewers and this was considered acceptable. A few discrepancies were resolved in discussion until mutual understanding of the inclusion criteria was improved and consensus was reached.

### Study inclusion

For the purposes of the present systematic map, searches included studies written in the English language that were published from 1^st^ January 2010 to 31^st^ December 2019. Observational (cross-sectional or cohort) studies that reported on the incidence and/or prevalence or mortality of selected infectious diseases affecting Ethiopian ruminants (cattle, sheep or goats) were considered for inclusion. The specific diseases to be studied were chosen based on experts’ opinion. Experimental studies, preliminary or pilot studies, studies that report on aggregated livestock data or outbreaks, studies which cover non-infectious diseases or other conditions (e.g. drought), as well as case report/series studies were excluded. Retrospective studies and surveys were also excluded. Narrative reviews, systematic reviews, maps and meta-analyses were included when primary data was available and these were recorded separately.

### Data collection process and items

After screening at full-text, two reviewers (TKT, ISMV) independently extracted bibliographical meta-data and study details from each study considered for inclusion into a pre-trialled extraction template. The template was piloted in 10 sample articles and all the discrepancies were discussed in detail by the reviewers. A coding scheme was developed iteratively during the systematic map screening process.

Bibliographical information extracted included the reference, year of publication, article, and journal title. Study details extracted included start and end date of data collection, experimental design, experimental factors, sampling method and statistical analysis. Study location details extracted included geographical region and agroecology. Population level details extracted included ruminant species, animal age (days, months and years and/or young, adult and old), breed, as well as the production system. The type of sample and the diagnostic method used, as well as the disease were also extracted. In addition, herd prevalence/incidence (%), individual prevalence/incidence (%) and mortality (%) were also extracted. Finally, a comments category allowed the entry of any potential reviewer’s comments to add further context to extracted information.

Primary data made available from published systematic reviews and meta-analyses were separately extracted and subsequently added to the main data set. In the coding scheme used, each line of the extraction form represented one prevalence value and not one article. For some categories multiple codes were applied, (for example, articles that reported results from more than one region, agroecology or experimental factor). If the relevant information could not be identified within the text, the field was left blank.

Upon completion of the data extraction, the dataset was harmonised and systematically checked for consistency by another author (LMD) to reduce entry errors. The extracted data were automatically cleaned using the statistical language R (https://www.r-project.org/).

### Systematic map data stored on Edinburgh DataShare

A copy of the Systematic map evidence base in the form of an excel file is stored on the Edinburgh DataShare platform (Tsouloufi *et al*., 2021). Thereby ensuring the long-term storage of the evidence base and allowing a persistent Digital Object Identifier (DOI) to be provided. As new searches are performed, the evidence base will be updated, therefore the data on Edinburgh DataShare represents a snapshot of the evidence base at the time of publication of this manuscript. The definitions of the categories and codes used in the systematic map are detailed in the meta-data of the excel file.

### Synthesis of the results

The evidence base identified within this systematic map is described in a narrative report and the descriptive statistics are presented in the form of figures. The evidence base is also visually presented as an interactive evidence map using the visualization platform Tableau (Washington, USA). The visualization includes multiple views of the dataset ranging from bibliographic information to study specific information targeted to specific users. In addition, the prevalence and mortality rates can also be explored (which are not reported on in this paper). The interactive evidence map can be accessed at https://www.livestockdata.org and aims to reduce time and resources required to retrieve evidence, thereby facilitating evidence based decision-making. The dashboard will be updated and act as a living systematic map of the ruminant disease evidence in Ethiopia. Updates will happen at least twice a year using a machine learning methodology applied by informatics experts (Goldfarb-Tarrant *et al*., 2020), a model used by other groups synthesising human health data (Shemilt *et al*., 2021).

### Study validity assessment

The validity of articles was not appraised as part of this systematic map in accordance with accepted systematic mapping methodological guidance (James *et al*., 2016). However, meta-data, such as study design, sampling method and experimental procedures, were extracted, allowing for internal and external validity of the included studies to be undertaken in future systematic reviews.

### Journal ranking

Along with the bibliographic information, a SCImago Journal Rank (SJR) indicator measurement was recorded. This is a relatively new, quantitative indicator metric that illustrates the journal’s scientific influence. It uses citation data from the Scopus database by calculating the average number of weighted citations received during a selected year for each document published in that journal during the previous three years. This indicator has been evaluated against the Thomson Reuters Scientific Impact Factor and was found to be a useful alternative for journal impact evaluation (Gonzalez-Pereira *et al*., 2010). This scheme categorises journals that are included in the Scopus database into quarters (Q1 to Q4, Q1 indicating journals with the highest prestige), while for a number of journals it is either not yet available (NA) or not indexed (NI). A single reviewer (ISMV) categorised the papers considered for inclusion according to their SJR indicators.

## Results

### Overall review descriptive statistics

The searches in the bibliographical databases resulted in 3164 records being collected from bibliographic databases and 66381 records were retrieved from search engines, specialist websites and organisational websites (Figure 1) in 2020. There were 1,456 articles in total considered eligible for full-text screening and data retrieval. Of the 1,456 articles screened, 716 were eligible for data extraction and subsequent inclusion in the systematic map database (Figure 1). The original endnote files were not archived for the bibliographic searches and therefore the searches were repeated in 2021. The searches in the bibliographical databases resulted in 3666 records in 2021, of which 3,073 were duplicates.

**Figure 1.**
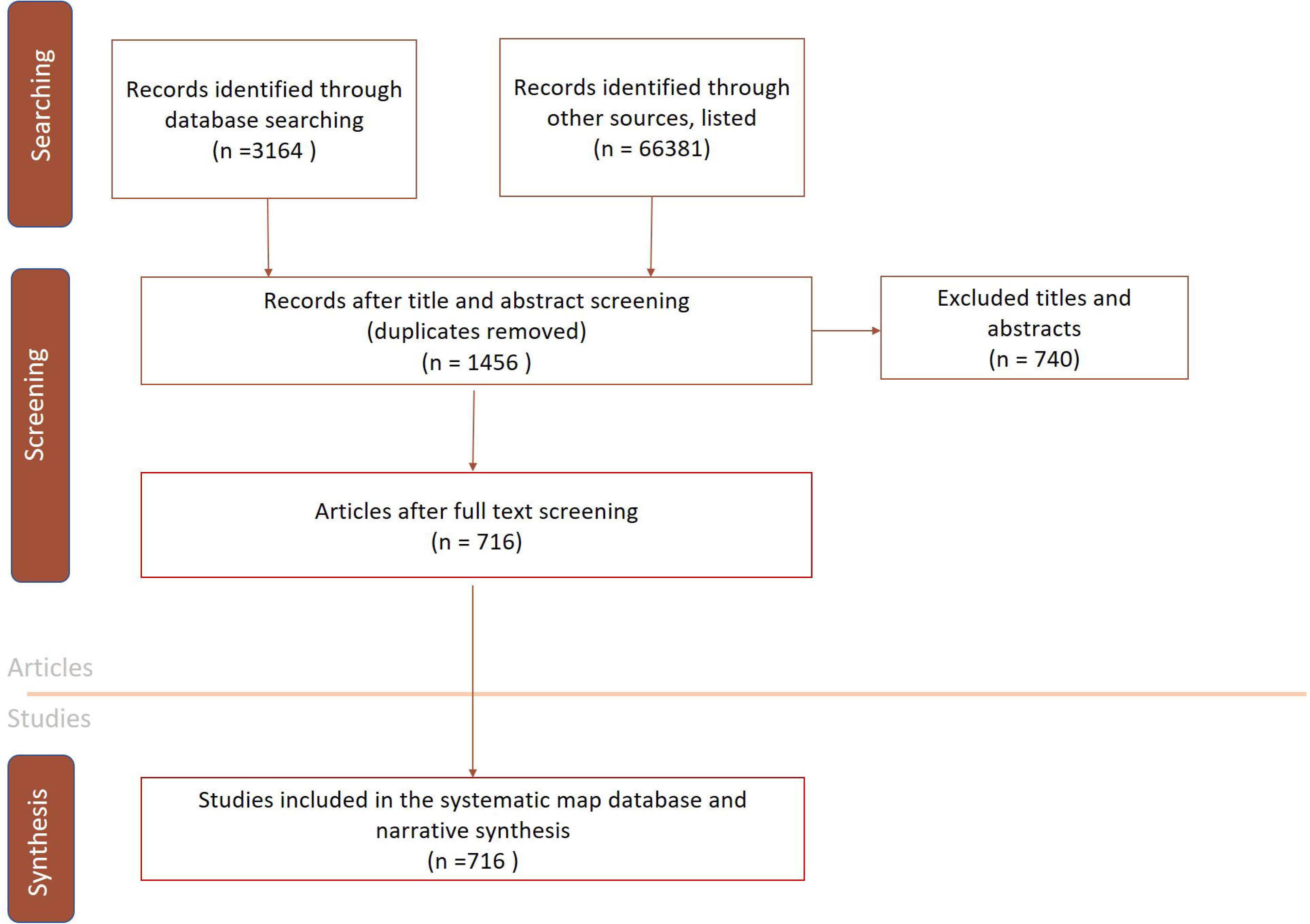
A conceptual diagram displaying the process of study selection and the respective number of studies included in the systematic map.

There were differences noted between the search results collected in 2020 and 2021 which are detailed in Supplementary File 1. These differences may be the result of bibliographic database changes during this time, but we also acknowledge the potential existence of human errors during the recording of search results. In 2021, a subset of searches were performed by a second author for validation.

### Bibliographical details and journal rankings

Journal articles were the most common type of publication (n= 696; 97.2%) followed by academic theses (n=17; 2.4%) and then conference proceedings (n=3; 0.23 %). Out of the 716 articles included in the map, 56 (7.8%) were published in a journal that ranked as Q1 in SCImago journal ranking system, 86 (12.0%) in a journal ranked as Q2, 24 (3.4%) in a Q3 journal and 34 (4.7%) in a Q4 journal. For 114 (15.9%) of the included papers the indicator was not calculated, while for 402 (56.1%) articles the indicator was not available as the respective journals were not included in the Scopus database (Figure 2). The journals chosen most often for publication were the *Journal of Veterinary Medicine and Animal Health* (NI) (n=65; 9.1%), followed by the *Ethiopian Veterinary Journal* (NI) (n=49; 6.8%) and *Tropical Animal Health and Production* (Q2) (n=47; 6.6%) as depicted in Figure 3.

**Figure 2.**
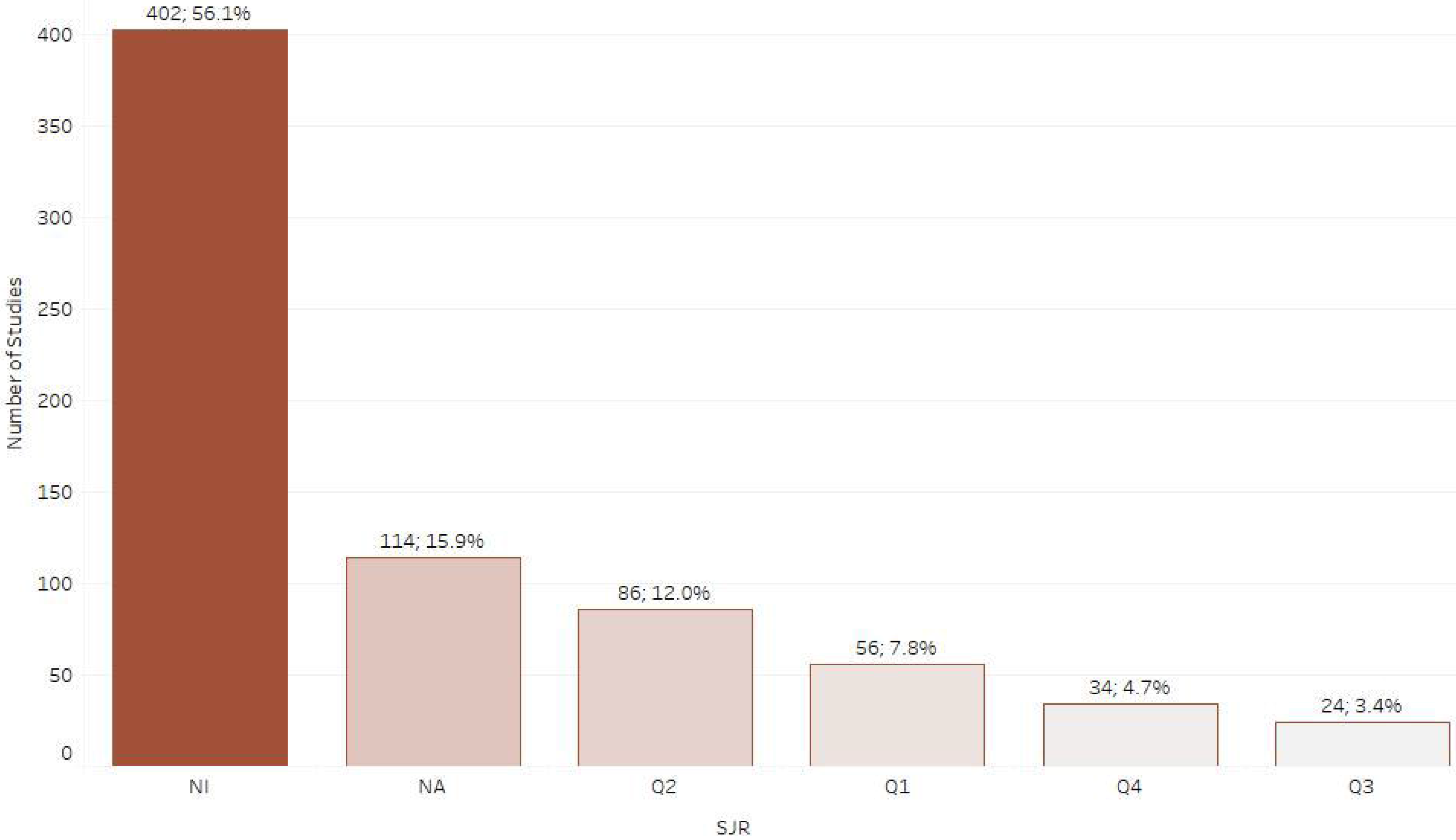
Number of retrieved studies categorised by SCImago journal ranking system (SJR).

**Figure 3.**
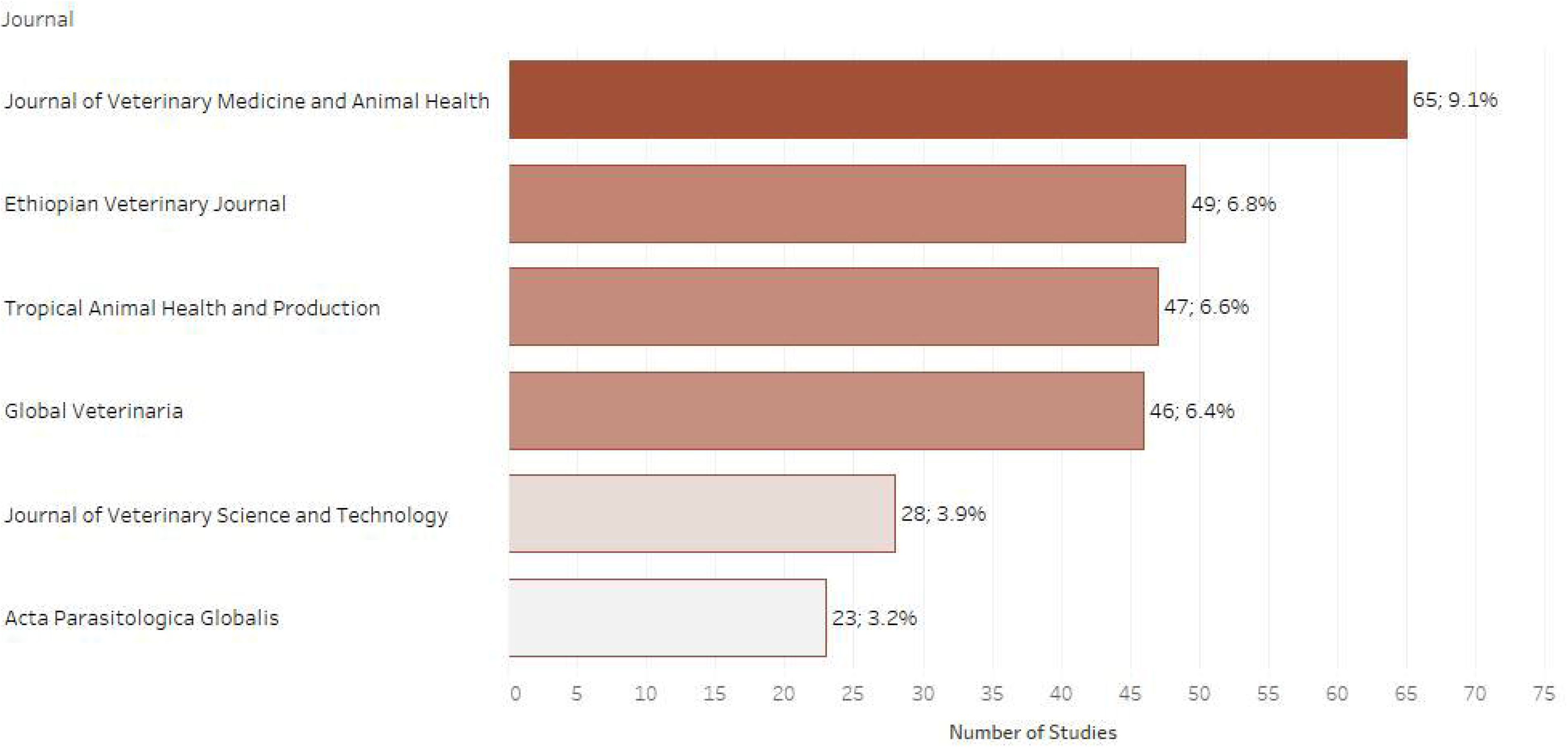
Number of retrieved studies published in each scientific journal (only journals with 20 or more retrieved studies are displayed).

### Distribution of studies per publication year

Figure 4 shows the number of studies published per year for the selected infectious diseases and their associated mortality in Ethiopia. Between the years 2012 and 2017 a considerable increase in the number of published studies was noted, with a peak of 101 and 97 studies observed in 2012 and 2017 respectively, when compared to 51 and 53 studies observed in 2010 and 2011 respectively. The number of published studies observed declined to 54 and 25 in 2018 and 2019 respectively.

**Figure 4.**
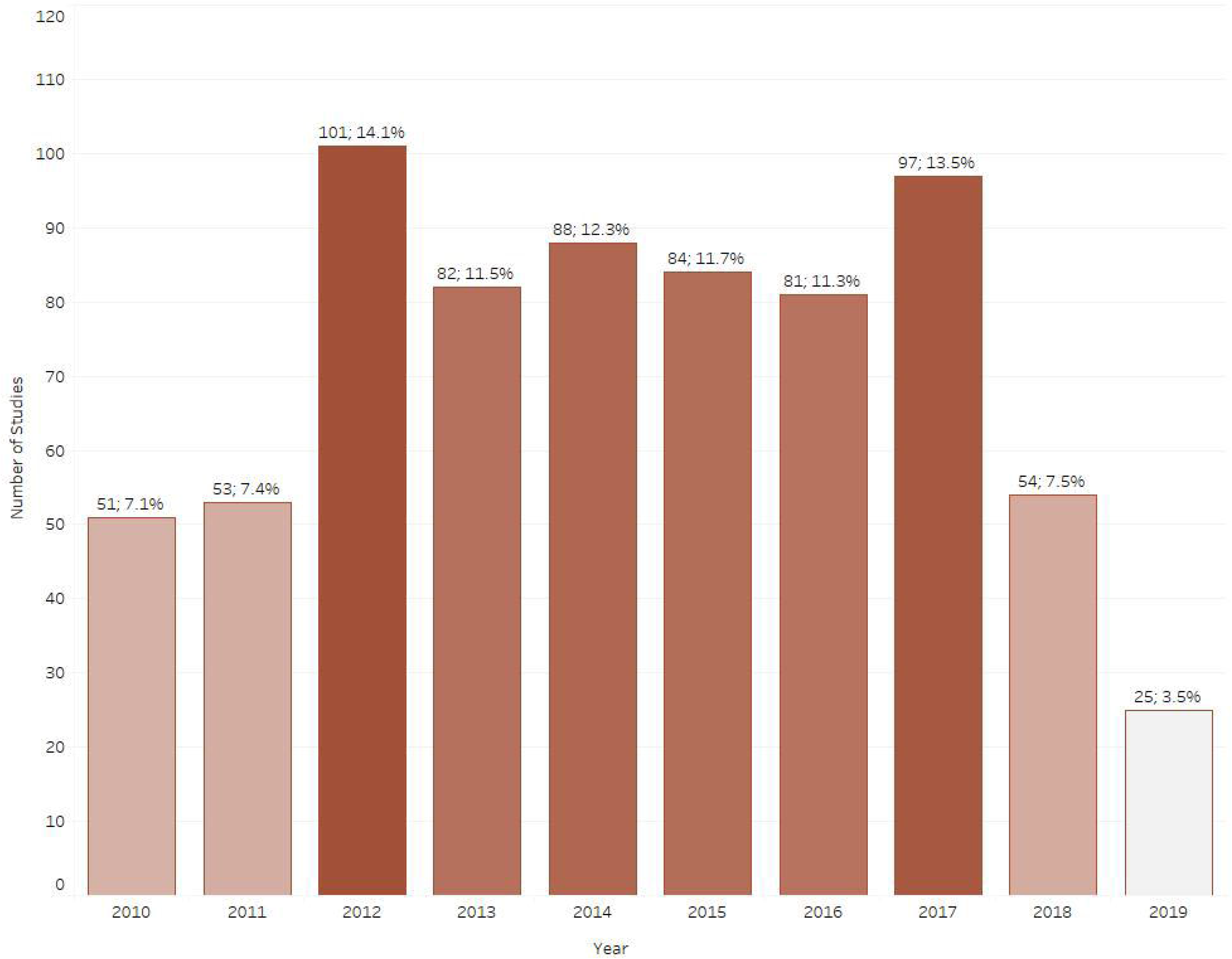
Number of retrieved studies published each year between 2010-2019

### Distribution of studies per geographical location

The distribution of published studies per disease and Ethiopian regional states is presented in Figure 5. Overall, most of the published studies (n=321; 44.8%) were conducted in Oromia, followed by Amhara (n=153; 21.4%) and the Southern Nations, Nationalities, and Peoples’ Region (n=128; 17.9%). The next most frequently studied regional states were Tigray (n=53; 7.4%), Addis Ababa (n=32; 4.5%) and Benishangul-Gumuz (n=31; 4.3%). The least studied Ethiopian regional states were Afar (n=23; 3.2%), Somali (n=17; 2.4%), the city of Dire Dawa (n=11; 1.5%), Gambella (n=7; 1.0%) and Harari (n=7; 1.0%). The most studied agroecological zone was the midland zone (n=398;5.6%), whereas lowland (n=158;22.1%) and highland (n=156;22.1%) zones were equally studied.

**Figure 5.**
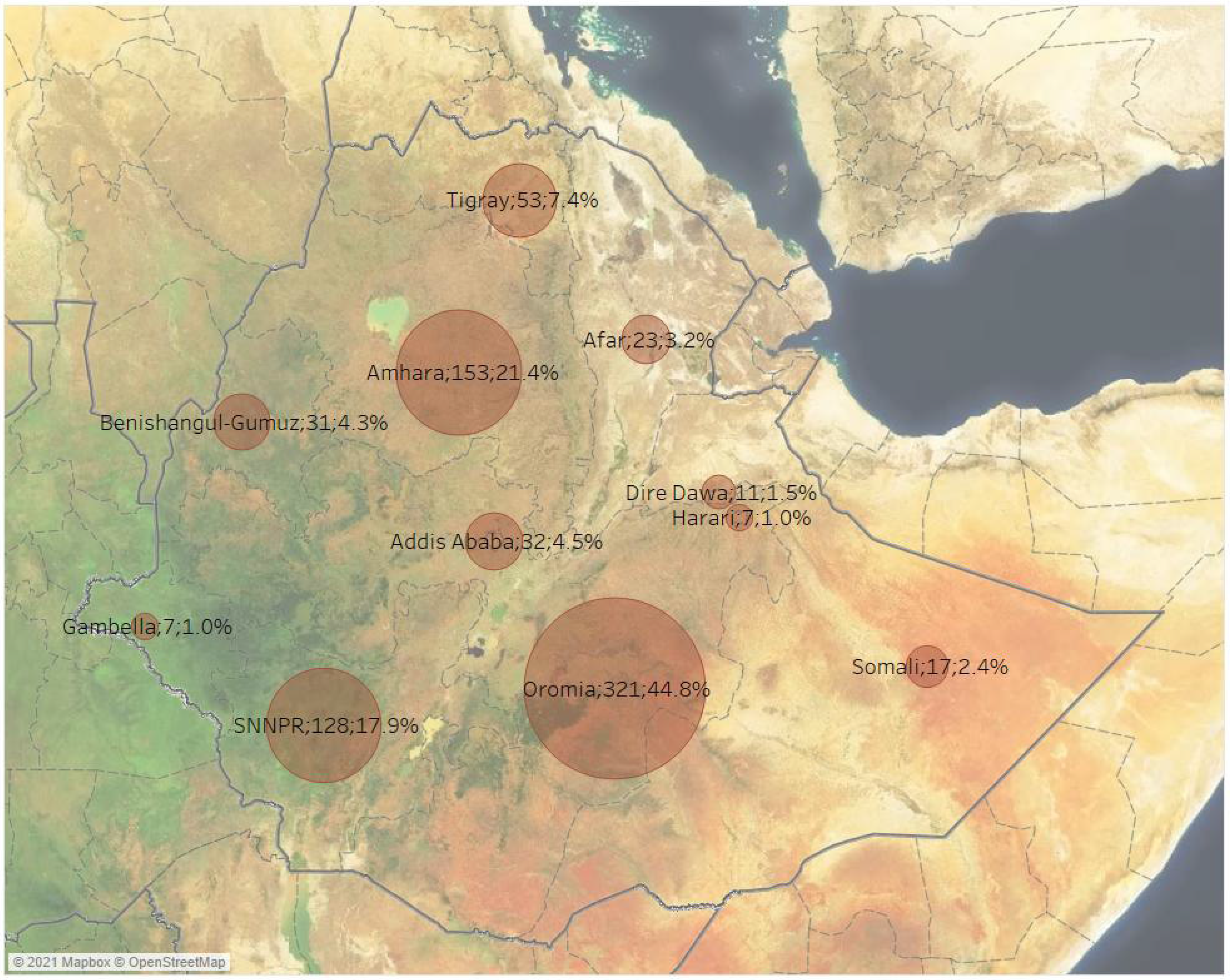
Number of retrieved studies by geographical region across Ethiopia (some studies included more than one geographical region)

### Distribution of studies per population and production system

Cattle (n=4469; 65.5%) were the most commonly studied population, followed by sheep (n=234; 32.7%) and then goats (n=203; 28.4%). Some studies reported their results as “small ruminants” (n=138; 19.3%). In almost half the studies animal breed was described (n=326;45.5%) and in certain cases the specific breed was mentioned such as Zebu, Boran or Holstein, however, more frequently generic categories were used such as local, cross, or exotic. There was considerable variation in the description of animal age amongst the studies, there are 128 (17.9%) studies in the dataset that have a description in terms of days, months and years, whereas a smaller number of studies (n=49;6.8%) are coded using the terms young, adult and old (the former coding took precedence to the later by the authors). There was considerable variation in terminology used to describe production systems and classifications such as extensive (156=n;21.8%), mixed (97=n;13.5%) or semi-intensive (40=n;5.6%) often being used.

### Distribution of studies per outcome

Regarding the outcomes measured, most of the included studies reported on individual disease prevalence (n=709; 98.3%), 65 studies (9.1%) reported on herd prevalence and 14 studies reported on mortality (2%).

### Distribution of studies per disease

Figure 6 shows the number of studies published for the selected infectious diseases and associated mortality in Ethiopia. Between 2010 and 2019, trypanosomosis was identified as the most studied disease in Ethiopia with 127 articles published overall (17.7%), followed by ectoparasite infestation (n=107; 14.9%), fasciolosis (n=87; 12.2%) and nematodiasis (n=87; 12.2%). The next most frequently studied diseases were echinococcosis (n=73; 10.2%), brucellosis (n=66; 9.2%) and bovine tuberculosis (n=53; 7.4%). Relatively low study numbers of studies were identified for cysticercosis (n=35; 4.9%), foot and mouth disease (n=26; 3.6%), contagious bovine pleuropneumonia (n=17; 2.4%), peste des petits ruminants (n=14; 2.0%), contagious caprine pleuropneumonia (n=14; 2.0%), pasteurellosis (n=14, 2.0%), dermatophilosis (n=11; 1.5%) and toxoplasmosis (n=11; 1.5%). For the selected publication years, infrequently studied ruminant diseases were lumpy skin disease (n=9; 1.3%), babesiosis (n=8; 1.1 %), sheep and goat pox (n=7; 1%) and cryptosporidiosis (n=7; 1.1%). Fewer than 5 articles were found for bovine anaplasmosis (n=5; 0.7%), bovine viral diarrhoea (n=3; 0.4%), cowdriosis (n=3; 0.4%), dermatophytosis (n=3; 0.4%), salmonellosis (n=3; 0.4%), bluetongue (n=3; 0.4%), rabies (n=3; 0.4%), Q fever (n=2; 0.3%), caseous lymphadenitis (n=2; 0.3%) and neosporosis (n=2; 0.3%). Only 1 article was found for each of the following diseases; blackleg, haemorrhagic septicemia, anthrax and infectious bovine rhinotracheitis. No evidence was found for the following ruminant diseases for the selected publication years when using our search strings: bovine genital campylobacteriosis, bovine spongiform encephalopathy, contagious agalactia, enzootic bovine leucosis, infectious necrotic hepatitis, leptospirosis, listeriosis, malignant catarrhal fever, ovine epididymitis, paratuberculosis, sarcocystosis, scrapie and trichomonosis.

**Figure 6.**
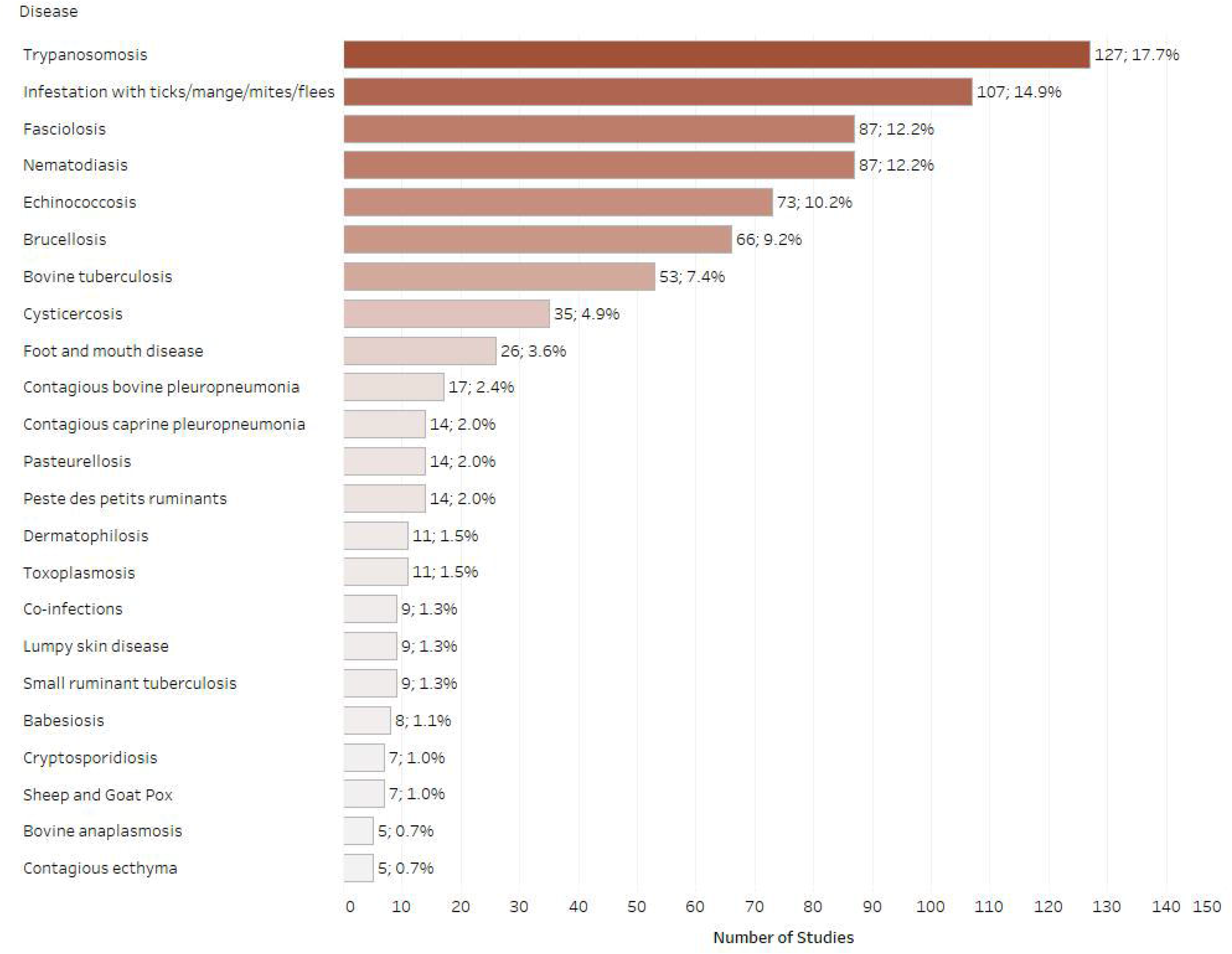
Number of retrieved studies published on each disease (only diseases with 5 or more retrieved studies are displayed, some studies included multiple diseases)

### Distribution of studies per diagnostic test

Figure 7 shows the number of studies published using a particular diagnostic test. In most of the included studies, the tests recommended by OIE were used for the detection of the respective disease agents. Specifically, most of the published studies used microscopy as a diagnostic test (n=324; 45.3%), followed by physical/post-mortem examination (PM examination) (n=146; 20.4%). There were 158 studies (22.1%) reporting to use serology techniques, covering a mixture of diagnostic tests (e.g. CFT, RBT, CIDT and ELISA). A small number of studies employed molecular tests or microbiological methods for the detection of the respective infectious agents. Frequently collected samples were serum (n=158;22.1%), whole blood (n=141;19.7%), carcass (n=128;17.9%) and faeces (n=116;16.2%).

**Figure 7.**
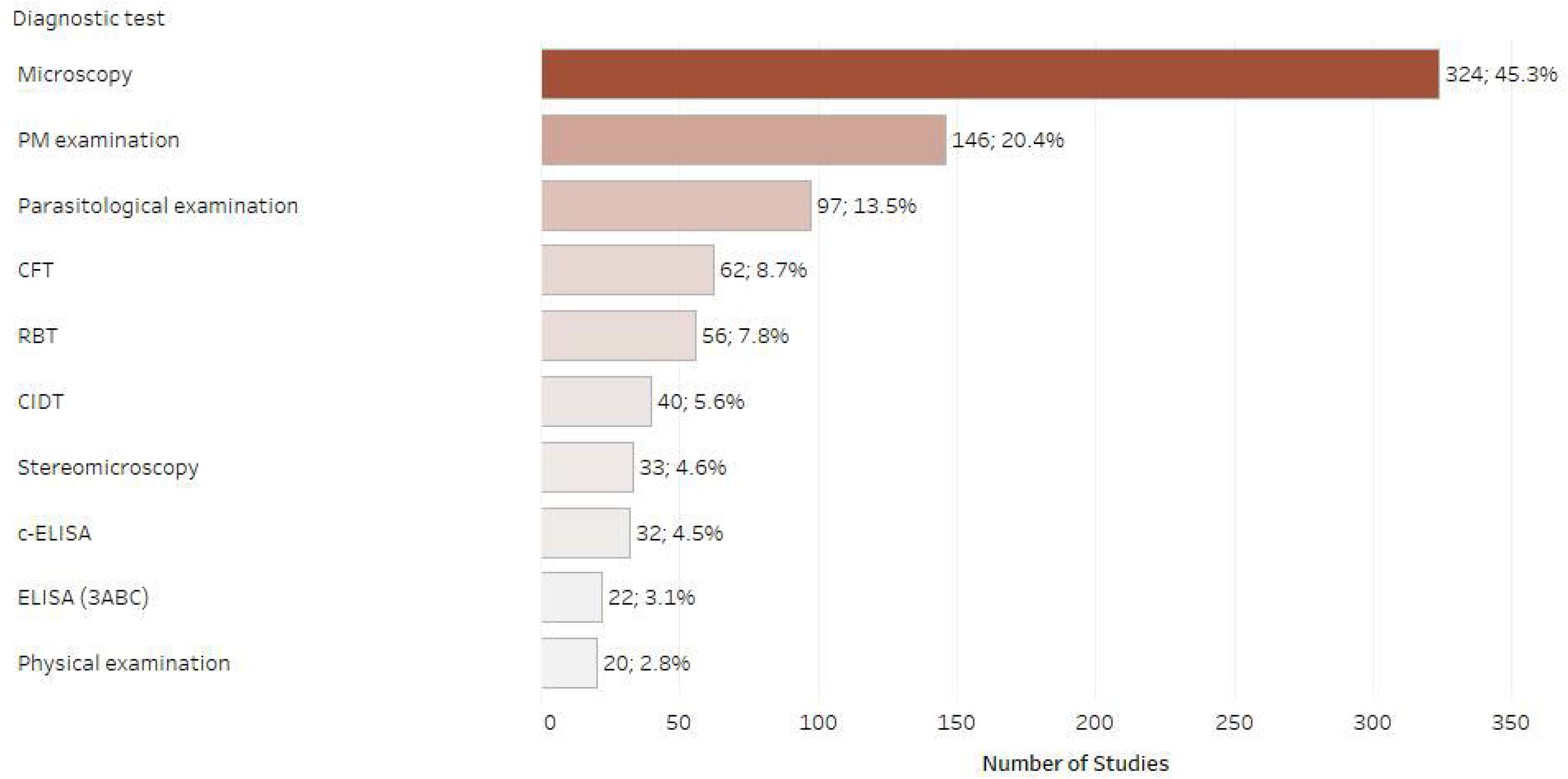
Number of retrieved studies published using each diagnostic test (only tests with 20 or more retrieved studies are displayed, studies frequently included multiple diagnostic tests).

### Distribution of studies per experimental factor

Figure 8 shows the most studied experimental factors. There was considerable variation in the use of terminology and factors studied. Age was the most commonly recorded factor (n=410; 57.3%), followed by sex (n=362; 50.6%) and then body condition score (n=257; 35.9%). Other commonly studied factors were geographical area, species, breed and management system.

**Figure 8.**
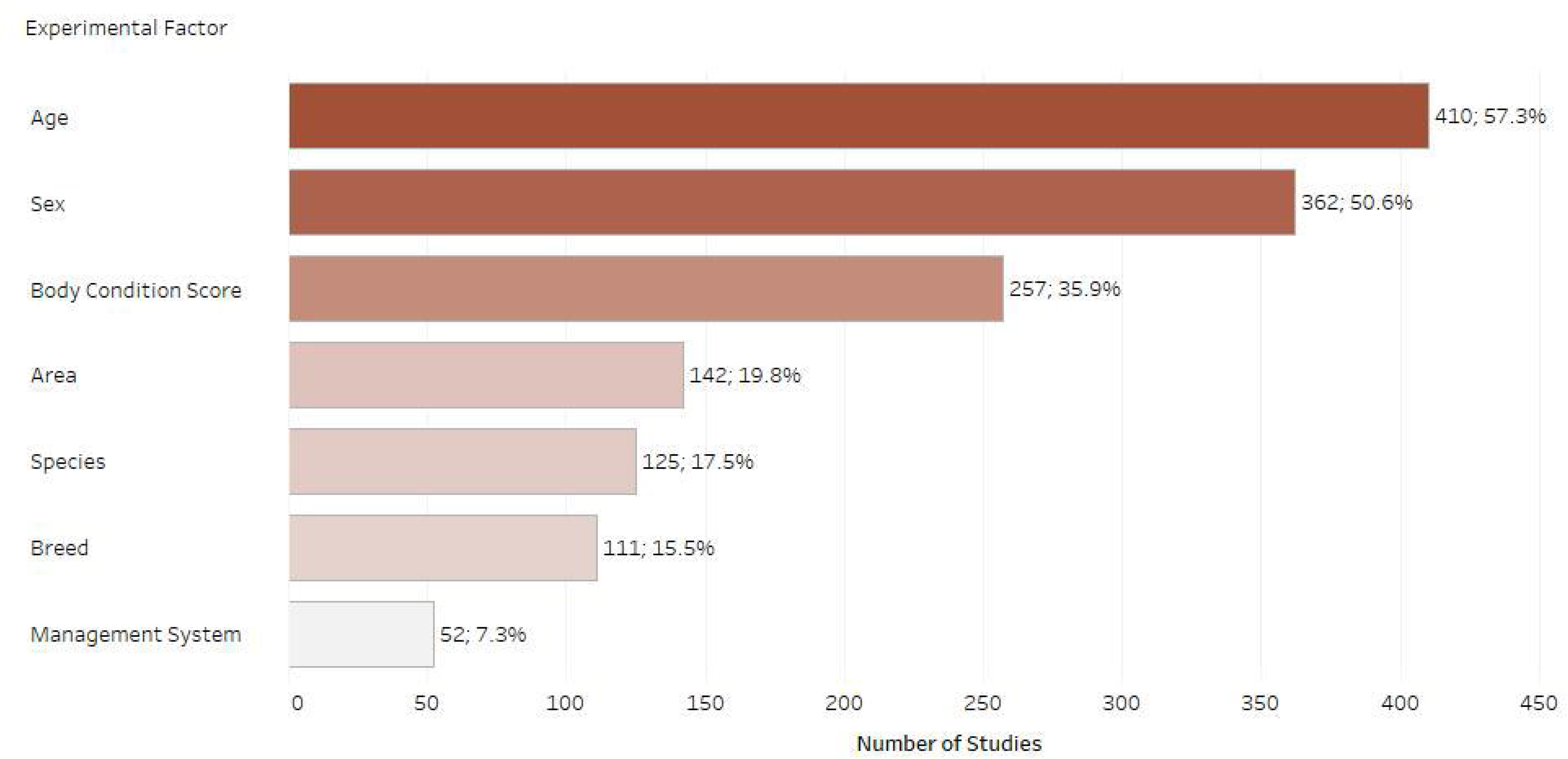
Number of retrieved studies published on each experimental factor (only experimental factors with 50 or more retrieved studies are displayed, studies frequently included multiple experimental factors).

### Distribution of studies per experimental design and sampling method

Most of the studies included in the present systematic map (n=697) were cross-sectional observational studies, while 5 were longitudinal observational studies. Most of the studies (n=642; 89.7%) reported on sample size. A considerable number of studies used a combination of sampling methodologies, with over half of the studies using simple random sampling (n=385; 53.8%).

## Discussion

### Summary of evidence

This review provides an up-to-date systematic map illustrating the available evidence for a diverse selection of ruminant diseases in Ethiopia between 2010 and 2019. Research funders and decision-makers have limited time and sometimes limited resources or knowledge to evaluate the existing evidence. A clear understanding and learning from previous efforts in the field of ruminant infectious diseases is necessary to prevent future redundant research. Hence, this map aimed to close a current synthesis gap by highlighting the knowledge gaps and clusters of this constantly growing research area. Most importantly, it should also improve the visibility of Ethiopian researchers by presenting research results from both published and grey literature.

Overall, primary research in ruminant infectious diseases seems to be relatively well reported in journal articles and the contribution of grey literature was not significant. Between 2010 and 2017, we noted a steady increase in the number of studies published for selected infectious diseases in Ethiopia (including two peaks in 2012 and 2017), followed by a sharp decline thereafter. The increase noted between 2012 to 2017 followed a publication momentum from 1996 to 2010, the number of veterinary articles being almost the same as those produced in Western Europe, North America, and the Pacific region when article output was normalized for GDP (Christopher and Marusic, 2013). According to the UNESCO institute for statistics (http://uis.unesco.org/), the gross domestic expenditures on research and development in Ethiopia peaked to 0.6% GDP in 2013 and sharply declined by about 50% in 2017. It is possible, therefore, that the decline in publications observed is due to the limited resources available. During the study period, there seems to be an uneven distribution of publications among the Ethiopian Regional States, with the vast majority concentrated in the most populous states, namely Oromia, SNNPR and Amhara. These states have the largest livestock density in the country, hosting 40% of the country’s livestock population (Leta and Mesela, 2014). Moreover, these states also concentrate the largest proportion of research institutes and veterinary schools in the country, which might account for the increased research output. Conversely, less populous Ethiopian states that rely heavily on extensive livestock production, such as Afar, Somali and Gambella, are under-represented in recent literature. All three states are considered Developing Regional States by the Ethiopian government due to the high poverty prevalence (www.unicef.org) and with the exception of Somali, have a considerably lower livestock population comprised mostly of small ruminants. This considerably fewer studies detected might be due to the mobility of the pastoral systems affecting accessibility, security, and resources.

For the purposes of this systematic map, we followed a comprehensive approach and we chose to include 52 infectious diseases that are known to affect ruminant productivity in LMICs. Our findings illustrate that the most researched diseases in the country for the selected period were trypanosomosis, ectoparasite infestations and nematodiasis, followed by fasciolosis, echinococcosis, brucellosis, and bovine tuberculosis. Regarding trypanosomosis, we identified over 100 studies (including 12 studies on small ruminant trypanosomosis), most of which were conducted in the western and south-western parts of the country (mostly in Oromia and Benishangul-Gumuz). This was an expected finding, as this area is known to be the major tsetse belt in Ethiopia (Balis and Bergeon, 1970). Our findings are partly confirmed by a recently published meta-analysis, which also identified a considerable number of publications on bovine trypanosomosis in Ethiopia from 1997 to 2015 (Leta *et al*., 2016). According to our findings, the increased research interest around trypanosomosis in the country contributed to the publication peak in 2012 and continued beyond 2015, with over 30 studies published between 2016 and 2019. Bovine trypanosomosis directly affects animal productivity and is implicated in increased mortality rates, incurring significant direct and indirect economic losses (Mattioli and Slingenbergh, 2013). In Ethiopia, there are a considerable number of veterinary parasitologists and projects e.g. Southern rift valley Tsetse Eradication Project (STEP) focused on the study of trypanosomosis (personal communication). Understandably, this topic has gathered a considerable research interest, which is reflected by the increase in number of publications. The most recent literature covers four *Trypanosoma* species (*T. vivax*, *T. congolense*, *T. brucei* and *T. evansi*) and the prevalence values reported vary greatly between the respective states and publication years, potentially reflecting different control and eradication efforts. Moreover, almost all the studies employed the micro-haematocrit centrifugation technique for the disease diagnosis, which is a simple and cost-effective diagnostic test that might also account for the high number of studies observed.

Infestation with ectoparasites is an important and often neglected condition in ruminants that leads to considerable economic losses due to impaired productivity, increased mortality and skin defects, while some ectoparasites (e.g. ticks) are vectors of important zoonotic agents. The most recent literature is evenly distributed across the publication years, with a publication peak noted in 2012, as with trypanosomosis. Most of the studies covering tick infestations in cattle, were conducted in the SNNPR regional state and reported highly variable prevalence values across the country. Nematodirases, in turn, seems to have been well-studied in Oromia and Amhara, mostly in the context of sheep gastrointestinal parasites; this is expected since the disease is considered one of the major constraints in the development of the Ethiopian livestock sector and was associated with 25% mortality and 3.8% weight loss in highland sheep (Bekele *et al*., 1992). Fasciolosis and echinococcosis are both widely prevalent in Ethiopia and incur significant economic losses due to the condemnation of the infected organs. Most of the published studies were conducted in Oromia and Amhara, which is an expected finding considering the number of abattoirs in these areas. Bovine brucellosis and tuberculosis, in turn, are important zoonoses that are widely prevalent in Ethiopia, and have a considerable economic impact due to animal productivity and market/trade impairments in urban and peri-urban areas. Both diseases are thought to follow a similar distribution within the country (Tschopp *et al*., 2013). This is reflected in the distribution of the recent literature as almost all regional states were represented (however there was low or no evidence from Harari, Dire Dawa, Gambella and Benishangul-Gumuz), while the majority of studies were conducted in Oromia, which is one of the highest milk producing areas in the country (Tschopp *et al*., 2013). Of note, we identified 12 publications on small ruminant tuberculosis, the majority of which were carried out in Afar pastoralist community and reported highly variable prevalence values.

Conversely, we identified limited evidence for specific infectious disease studies over the last decade, which are listed as important by OIE. Interestingly, we identified only 14 studies on PPR during the course of this review, eight of which were conducted after 2015, when the PPR Global Control and Eradication strategy was launched (OIE, 2016). Most of the studies were conducted in pastoral areas (e.g. Afar, Gambella) and suggest extensive circulation of PPR virus among the small ruminant population. PPR is a disease of considerable economic importance due to its high morbidity and mortality and is endemic in most pastoral areas of Ethiopia. A new PPR elimination program is currently underway in pastoralist areas; however, the currently insufficient understanding of PPR epidemiology in the country is likely to undermine future elimination or vaccination efforts. An effective local vaccine roll-out strategy, in turn, might account for the limited research interest observed for diseases, such as black leg, haemorrhagic septicaemia, sheep and goat pox, CBPP and CCPP. With regard to CCPP, a published meta-analysis highlighted the lack of data for this particular disease (Asmare *et al*., 2016); the research interest in this disease seems to be declining recently, as we identified only five studies published between 2015 and 2019. The evidence for CBPP follows a reverse pattern, with most studies published between 2015 and 2019; even though there is a paucity of research in the country, the disease is widespread and considered to be one of the most important impediments to livestock development in Ethiopia (Abdela and Yune, 2017). Moreover, we found very few studies investigating vector-borne diseases, excluding trypanosomosis (e.g. piroplasmosis, cowdriosis, LSD, bluetongue), which was somewhat expected considering the generally limited research evidence available for these diseases (Asmare *et al*., 2017). It was also noted that there is a considerable lack of research on the pathogens implicated in calf diarrhoeas (e.g. BVD, *Escherichia coli, Salmonella* spp., and *Cryptosporidium* spp.), which is the most common cause of neonatal calf mortality. In Ethiopia, calf morbidity and mortality were ranked as the second biggest issues for dairy production in Ethiopia after mastitis (ILCA, 1994). Finally, diseases like IBR and BVD that are prevalent throughout the country seem to be less studied, potentially due to the lack of diagnostic resources.

Most importantly, a knowledge gap was identified for some diseases including anthrax, enzootic bovine leucosis, leptospirosis, listeriosis, contagious agalactia, enterotoxaemia or paratuberculosis. This was not surprising considering that most of these diseases are rarely occurring (e.g. listeriosis, enzootic bovine leucosis), and are underdiagnosed (e.g. contagious agalactia) or typically documented during large outbreaks (e.g. anthrax). The complete absence of data on leptospirosis over the last decade is deemed a major limitation of the current evidence base, however, as it is a globally important zoonotic disease and the available prevalence/incidence studies in Ethiopia are rather outdated (de Vries *et al*., 2014)

The diagnostic tests used in epidemiological studies have a major impact on reported prevalence and introduce specific limitations to the understanding of the current evidence base. According to our findings, most of the studies employed tests recommended by the World Organization for Animal Health (2012). Although the tests varied according to their respective disease, the overall most used diagnostic tests were microscopy and serology followed by physical examination/post-mortem diagnosis (i.e. abattoirs), whereas a very small number of studies employed molecular or microbiological methods. Microscopy is a simple and cost-effective diagnostic method; however, despite its high specificity, it has low analytical sensitivity, particularly for parasitic diseases, due to variable levels of parasitaemia. Cross-sectional studies, in turn, that employ serological tests, are unable to conclude on the true prevalence, as antibody levels can fluctuate, especially during the early course of infection. The utility of antibody tests should be further considered, particularly in the context of resource limiting settings, as cattle are managed poorly under chronic malnutrition, high burden of gastrointestinal parasites and concurrent infections, which could potentially affect their immune responses. Hence, more sensitive techniques including isolation and molecular detection of agents need to become more accessible for rapid diagnosis, to improve the accuracy of prevalence estimates. Finally, in our review we identified studies that were conducted in abattoirs, mostly covering cattle echinococcosis and fasciolosis in Oromia, SNNPR and Amhara. While this is a cost-effective option for an epidemiological study that enables access to larger sample sizes, it is prone to non-random error and information biases (e.g. poor record keeping) that can affect the prevalence estimates (Delgado-Rodriguez and Llorca, 2004).

Regarding disease-associated mortality, our review findings indicate that data are remarkably scarce for Ethiopia and are mostly in the context of surveys or outbreak investigations. The studies identified reported on mortality in Oromia, SNNPR, Amhara and Tigray, but age was often not recorded, however young stock mortality is generally considered high in Sub-Saharan Africa and approximately ranges from 9% to 45% (Wymann *et al*., 2006; Bolajoko *et al*., 2020; Wong *et al*., 2021). The reported mortality percentages ranged from 1 to 25% for studies published from 2010 to 2019, although no clear definition of the measure of mortality was provided. This is an unsurprising finding, as measuring ruminant mortality in LMICs can prove challenging due to the significant under-reporting of livestock losses and the limited baseline data in livestock numbers (Catley *et al*., 2014; Wong *et al*., 2021). Moreover, mortality rates are higher when reported in farmer surveys; this might be attributed to the assignment of livestock culling or sales in this category by the respondents or the underestimation of other concurrent factors that can cause livestock mortality, such as poor nutrition, mycotoxins etc. Determining a measure of mortality can be challenging (Wong *et al*., 2021); future studies focussing on this aspect should include information about the measure of mortality (mortality rate, mortality risk or other) and adequate contextual data, such as study design, sampling method, sample sizes etc., which will aid the interpretation of the findings.

### Limitations of the evidence base

A limiting factor to the current evidence base is the lack of established ontologies or agreed definitions with regards to specific data categories such age, mortality and production systems, which limits the comparability of results across studies. Furthermore, there is specific information that was occasionally not reported in the compiled studies, such as the descriptive statistics of the included population, the validity or performance characteristics of each diagnostic assay used and the standards followed for each assay or the vaccination status of the animals when serology was used. All these concur with the already recognised issue of reporting inadequacies in veterinary observational studies and the need for wider application of reporting standards (e.g. STROBE) (Sargeant and Connor, 2014).

### Limitations of the review

This systematic mapping review bears limitations that should be considered for future iterations. Firstly, a few deviations from the original protocol (Tsouloufi *et al*., 2020) were made, mostly due to time and resource constraints. Secondly, books and monographs were not accessed for the purposes of the present study, and no snowballing technique was applied to retrieve extra references; therefore, there is a possibility that a small number of resources were inadvertently missed. Secondly, this study aimed to explore the evidence with specific date and language limitations, which may have affected the final distribution of the evidence. Specifically, the searches were limited to English language and it is possible that manuscripts written in the Amharic or other local languages were missed. However, although these are expected to represent a small percentage of the published literature, the inclusion of non-English language databases would be a worthwhile addition for any future updates. Similarly, as we focused on publication years 2010 to 2019 and no updated searches have yet been performed, this review fails to cover any research performed between then and the publication of this paper. Thirdly, although the review follows a comprehensive approach and covers 52 infectious diseases, there is a possibility that some conditions were omitted. Fourthly, we only studied disease prevalence at a regional level; future iterations should consider study status on a district level to fully understand the epidemiology of the problem. Although we described the distribution of papers according to their SCImago ranking, this was not intended to be a comprehensive bibliometric study and thus, no definitive conclusions should be drawn in these respects. Finally, the original Endnote files for the bibliographic searches were not archived, which has resulted in the paper not reporting on all the intermediate steps of the screening process as recommended by the guidelines (Haddaway *et al*., 2017) e.g. duplicate removal, records after title screening etc.

### Review conclusions and recommendations

This systematic map provides a first comprehensive synthesis of recent evidence available on ruminant infectious disease frequency and disease-associated mortality in Ethiopia. Overall, the published research seems concentrated on a few infectious diseases of parasitic and bacterial origin, whereas most of the selected diseases are either under-represented or absent. This review is intended to form the basis for further primary research by identifying key knowledge gaps and be a starting point for the synthesis of information in focused systematic reviews. As the intention is to keep this map an Open Access source, interested stakeholders will have the opportunity to view study data with unlimited accessibility, the lack of which is often mentioned to hamper evidence use (Haddaway *et al*., 2016). The work underway on automating the process will enable evidence to be represented as living map accessible form the online visualisation (Goldfarb-Tarrant *et al*., 2020).

### Implications and recommendations for policy and research

Future primary research efforts should focus on addressing the aforementioned limitations of the evidence base. Specifically, when conducting an observational study, efforts should be made to adhere to specific reporting standards (e.g. STROBE), so that internal and external validity can be adequately evaluated. On these grounds, it is vital that consensus statements are released on livestock ontologies, particularly in areas such as animal age or mortality, as this will ensure the interoperability of the reported results. Future studies should also focus on consolidating evidence from geographically distinct literature (i.e. states such as Afar, Somali and Gambella which have been underrepresented over the last decade) and local languages to ensure that local knowledge has an appropriate coverage. At this point, it should be noted that the number of publications for a selected diseases is a measure of research productivity and does not necessarily correlate with the burden or impact of illness in the country; this map should therefore not be consulted solely for the disease prioritization in the country. Some significant knowledge gaps in the country were identified regarding the frequency of some diseases and are presented in Table.1.

**Table 1.** Gaps in knowledge of certain diseases

| **Disease** | **Gaps in knowledge** |
| --- | --- |
| **PPR** | Targeted for eradication by 2030; limited evidence collected from most states, no recent evidence for some Ethiopian states (e.g. Afar and Gambella) |
| **CCPP and CBPP** | Some Ethiopian states (e.g. Amhara) have no collected evidence or have no recent evidence (e.g. Afar) |
| **Q fever** | Emerging zoonotic disease in LMICs, with significant socio-economic implications; only two studies identified |
| **Leptospirosis** | Significant zoonotic disease with generally limited and dated evidence, no study in the last decade identified |
| **Vector-borne diseases excluding trypanosomosis (e.g. piroplasmosis, cowdriosis, bluetongue and dermatophilosis)** | Well-recognised causes of compromised productive and reproductive performance of small ruminants in SSA; a dearth of published evidence over the last decade |
| **Small ruminant brucellosis** | Compared to bovine brucellosis, a dearth of studies discusses small ruminant brucellosis recently |
| **Diarrhoea-inducing infectious agents (e.g. BVDV, *Salmonella* spp, *Cryptosporidium* spp, *Escherichia coli*)** | Despite the importance of the condition in cattle production, very little is known about its status |
| **Disease co-infections** | Their role in livestock productivity is commonly underestimated, as specific diseases are usually targeted |
| **Disease-associated mortality** | Current knowledge comes exclusively from farmer surveys; best measure of mortality needs to be decided |
| **Paratuberculosis** | Only case reports and limited prevalence studies are available in SSA |

### Implications for future evidence synthesis

The map highlights areas where there may be sufficient data to justify a systematic review. We would strongly advise the inclusion of stakeholder engagement for the formulation and prioritization of the research question to ensure the relevance of the intended outputs. Furthermore, the present systematic mapping review will be updated, to ensure new evidence is included in a timely manner. Areas for future evidence synthesis have been highlighted by this map, in some cases they have been addressed by recent publications.

i. The results of a meta-analysis on trypanosomosis published in 2016 could be updated as 60 additional studies were published up to the end of 2019 (Leta *et al*., 2016).
ii. The results of a meta-analysis on brucellosis conducted in 2014 has been updated (Tesfaye *et al*., 2014).
iii. Although tick infestations are mostly covered in a recent systematic review 2017, evidence is constantly growing, but there is no recent systematic review covering all ectoparasites (Asmare *al.,* 2017).
iv. No recent systematic review or meta-analysis covering different endoparasites has been published.

## Acknowledgements

The authors would like to thank Prof Alemayehu Lemma (College of Veterinary Medicine and Agriculture, Addis Ababa University) for his comments on the manuscript.

